# Nested PCR to optimize *rpoB* metabarcoding for low-concentration and host-associated bacterial DNA

**DOI:** 10.1101/2025.03.17.643641

**Authors:** Laëtitia Leclerc, Gabryelle Agoutin, Thierry Brévault, Antony Champion, Jean Mainguy, Nicolas Nègre, Sudeeptha Yainna, Géraldine Pascal, Sophie Gaudriault, Jean-Claude Ogier

**Affiliations:** DGIMI, Univ Montpellier, INRAE, Montpellier, France; GenPhySE, Université de Toulouse, INRAE, ENVT, 31326, Castanet Tolosan, France; AIDA, Univ Montpellier, CIRAD, Montpellier, France; DIADE, Univ Montpellier, IRD, Montpellier, France; Université de Toulouse, INRAE, BioinfOmics, GenoToul Bioinformatics facility, 31326, Castanet-Tolosan, France; Université de Toulouse, INRAE, UR 875 MIAT, 31326, Castanet-Tolosan, France

**Keywords:** metabarcoding, host-associated microbiota, *rpoB*, nested PCR, insect

## Abstract

**Background:** Housekeeping genes have proved effective taxonomic markers for characterizing bacterial microbiota in short-read amplicon metabarcoding studies. A region of the *rpoB* gene, in particular, has been shown to minimize OTU overestimation bias with a high degree of accuracy, providing better species-level taxonomic resolution (Ogier et al. BMC Microbiol. 19:171). However, the primers for *rpoB* are highly degenerate, leading to potential problems in the amplification of bacterial DNA present at low concentration in the sample or embedded within eukaryotic matrices, as for the host-associated microbiota. We addressed these limitations by using a two-step PCR approach to optimize the *rpoB* procedure. The first PCR amplifies a 906-nucleotide region of the *rpoB* gene with the classical primers, referred to here as outer primers, and the second PCR then uses primers incorporating Illumina adapters, referred to here as inner primers, to amplify a 435-nucleotide subregion, the taxonomic marker for metabarcoding.

**Results:** We first used *in silico* approaches to evaluate the universality of the outer and inner *rpoB* primers. We then tested the nested *rpoB* PCR method on commercial mock samples of known composition. The nested PCR approach increased amplification efficiency for dilute samples without biasing the bacterial composition of the mock sample revealed by metabarcoding relative to single-step PCR. We also tested the nested *rpoB* PCR method on field-collected samples of the lepidopteran *Spodoptera frugiperda*. The nested PCR outperformed single-step PCR, increasing amplification efficiency for bacterial DNA present at low concentrations (oral secretions from *S. frugiperda*) or embedded in eukaryotic DNA matrices (*S. frugiperda* larvae).

**Conclusions:** This method provides a promising new strategy for characterizing insect-associated microbiota that can also be applied to other host microbiomes.

## Background

Microbial communities are frequently characterized by polymerase chain reaction (PCR) amplification followed by next-generation sequencing (NGS), particularly through short-read metabarcoding approaches, typically targeting variable regions of the 16S rRNA gene, such as the V3-V4 regions. However, short-amplicon sequencing studies based on the 16S rRNA gene are subject to certain limitations, particularly in terms of taxonomic resolution, which is often restricted to the genus and family levels [1, 2]. Full-length 16S rRNA gene sequencing with PacBio technology has been shown to improve taxonomic resolution in human microbiome samples [3], but its use is currently limited due to the higher cost required to obtain an equivalent number of reads per sample.

Housekeeping genes encoding conserved proteins ubiquitous in the bacterial kingdom provide a possible alternative to the short amplicon of the 16S rRNA gene. These genes evolve much more rapidly than the 16S rRNA gene and can therefore be used to differentiate between bacterial lineages [4]. Moreover, unlike the 16S rRNA gene, housekeeping genes are generally present as a single copy in genomes, minimizing biases in diversity assessment and providing a more accurate representation of bacterial community composition. Housekeeping genes have proved effective for the characterization of bacterial diversity, particularly at the species level, across various ecosystems [4–8]. The *rpoB* marker, in particular, has been successfully used in metabarcoding analyses of samples rich in Pseudomonadota (formerly Proteobacteria), such as entomopathogenic nematodes, and has been shown to have a finer taxonomic resolution than the 16S (V3V4) marker [5, 9]. In particular, phylogenies based on the same *rpoB* subregion are highly concordant with core genome phylogenies [10], indicating that this subregion is a robust marker for precise bacterial identification.

One key challenge in the metabarcoding of microbial communities is detecting prokaryotic DNA in samples with very low bacterial DNA concentrations, or in samples in which eukaryotic DNA predominates, as in the host-associated microbiota. In most cases, physically separating microbes from host tissues is difficult or even impossible, making global DNA extraction the only option. However, global DNA extraction typically results in a low microbial-to-host DNA ratio, limiting the efficiency of amplification with bacterial primers. This issue is further exacerbated by the use of primers targeting housekeeping genes, such as *rpoB*, which has a highly variable sequence, necessitating the use of degenerate bases to maximize species coverage. This may increase cross-reactivity with eukaryotic DNA, further hampering target amplification. Finally, in metabarcoding protocols, the long sequence and GC-rich regions of barcoding primers (about 60 nucleotides, including the target region, index sequences, and Illumina adapters) reduce the specificity of hybridization, making it even more challenging to amplify bacterial DNA from a host DNA matrix.

Nested PCR can be used to increase amplification sensitivity. In nested PCR approaches, there are two successive reactions, the first to amplify a larger region, and the second to amplify a smaller region nested within the first amplicon. Nested PCR is particularly useful for samples with low target DNA concentrations [11], but this method has only rarely been used in microbial ecology [12], and its potential for metabarcoding has yet to be explored.

In this study, we evaluated the potential of a nested PCR approach combined with a metabarcoding analysis with *rpoB* primers for characterizing bacterial communities in samples with low bacterial DNA concentrations or in which the bacterial DNA is embedded in a eukaryotic matrix. For validation of the efficacy of this strategy, we compared the performance of the single-step and nested *rpoB* PCR methods on a set of mock samples. We then compared the efficiency of the nested *rpoB* PCR approach to that of single-step PCR on biological samples consisting of oral secretions (OS) from the lepidopteran *Spodoptera frugiperda*, which contain low concentrations of bacterial DNA, and *S. frugiperda* larvae in which the bacterial DNA is diluted in large amounts of eukaryotic DNA.

## Materials and Methods

### Biological material

The two commercial mock samples contained DNA from eight bacterial strains in equal proportions (Mock_8sp; ZymoBIOMICS Microbial Community DNA Standard; D6305), and various proportions DNA from the same eukaryotic species (Mock_8sp log; ZymoBIOMICS Microbial Community DNA Standard II_log distribution; D6311). The composition of the mock communities is described in Table 1.

**Table 1.** Composition of the commercial DNA mock samples.

**Mock\_8sp**
| Species | Expected composition (%) |  |  |  |
| --- | --- | --- | --- | --- |
|  | Genomic DNA | 16S copies | Genome copies | Cell numbers |
| *Pseudomonas aeruginosa* | 12 | 4.2 | 6.1 | 6.1 |
| *Escherichia coli* | 12 | 10.1 | 8.5 | 8.5 |
| *Salmonella enterica* | 12 | 10.4 | 8.7 | 8.8 |
| *Limosilactobacillus fermentum* | 12 | 18.4 | 21.6 | 21.9 |
| *Enterococcus faecalis* | 12 | 9.9 | 14.6 | 14.6 |
| *Staphylococcus aureus* | 12 | 15.5 | 15.2 | 15.3 |
| *Listeria monocytogenes* | 12 | 14.1 | 13.9 | 13.9 |
| *Bacillus spizizenii* | 12 | 17.4 | 10.3 | 10.3 |

**Mock\_8sp log**
| Species | Expected composition (%) |  |  |  |
| --- | --- | --- | --- | --- |
|  | Genomic DNA | 16S copies | Genome copies | Cell numbers |
| *Pseudomonas aeruginosa* | 89.1 | 95.9 | 94.8 | 94.9 |
| *Escherichia coli* | 8.9 | 2.8 | 4.2 | 4.2 |
| *Salmonella enterica* | 0.89 | 1.2 | 0.7 | 0.7 |
| *Limosilactobacillus fermentum* | 0.89 | NA | 0.23 | 0.12 |
| *Enterococcus faecalis* | 0.089 | 0.069 | 0.058 | 0.058 |
| *Staphylococcus aureus* | 0.089 | 0.07 | 0.059 | 0.059 |
| *Listeria monocytogenes* | 0.0089 | 0.012 | 0.015 | 0.015 |
| *Bacillus spizizenii* | 0.00089 | 0.00067 | 0.001 | 0.001 |

Wild *S. frugiperda* larvae and their oral secretions (OS) were collected from maize (*Zea mays*, Poaeceae) fields near Velingara in the Anambé Valley in Southern Senegal (13.163268/-14.039137) in September 2021. We collected OS from approximately 100 larvae with flexible forceps that had been disinfected with 70% ethanol, and sterile plastic Pasteur pipettes. The samples were pooled to obtain a final volume of 200 µL and stored at approximately 4°C for a few hours before being frozen at −20°C.

### Extraction of bacterial DNA from insect samples

Total DNA was extracted from both the OS pool (about 200 µL) and six individual larvae (posterior half body): three biological replicates at stage L3 and three at stage L5. OS and larval samples were lysed by freezing at −80°C for 15 min and then heating at 80°C for 20 min. We then added 100 μL (500 µL for the larval samples) Quick Extract lysis solution (Bacterial DNA extraction kit from Epi-centre, USA), 1 μL (2 µL for the larval samples) Ready-Lyse Lysozyme Solution (Epi-centre, USA) and 20 µL (50µL for the larval samples) EDTA (0.5 M, pH 8). The samples were mechanically lysed with six 2 mm glass beads and three cycles of 7 m/s for 40 s in a FastPrep device (MP Biomedicals, Illkirch-Graffenstaden, France). The complete lysis of insect and prokaryotic cells was achieved by incubation at room temperature for 90 min and then at 55°C with 10 µL (20 µL for the larval samples) proteinase K (20 mg/mL), with shaking at 1200 rpm until the solution cleared. RNA was eliminated by incubation with 10 μL (20 µL for the larval samples) RNase A at 20 mg/mL (Invitrogen PureLinkTM RNaseA, France) for 15 min at 37°C. After a final incubation with 100 µL protein precipitation solution (Wizard kit, Promega, France) for 5 min on ice, the samples were centrifuged for 5 min at 17,000 x *g* and 4°C and the supernatant was recovered. OS samples were subjected to phenol and chloroform treatments due to the presence of polysaccharides. The DNA present in the OS and larval samples was precipitated with absolute ethanol and washed with 70% ethanol. The extracted DNA was resuspended in 50 μL ultrapure water and stored at −20°C. We assessed the levels of contaminating bacterial DNA at each of the various steps in DNA sample preparation, by including several negative control extractions with sterile ultrapure water.

### Library preparation and Illumina MiSeq sequencing

We used two PCR strategies to generate *rpoB* amplicon libraries for MiSeq sequencing: single-step PCR, involving direct amplification, and nested PCR, consisting of two successive PCRs (Fig. 1). Each PCR was performed in duplicate (mock samples) or triplicate (OS samples). Our protocol also included RNase free water as a negative control and positive *E. mundtii* genomic DNA as a positive control.

**Figure 1.**
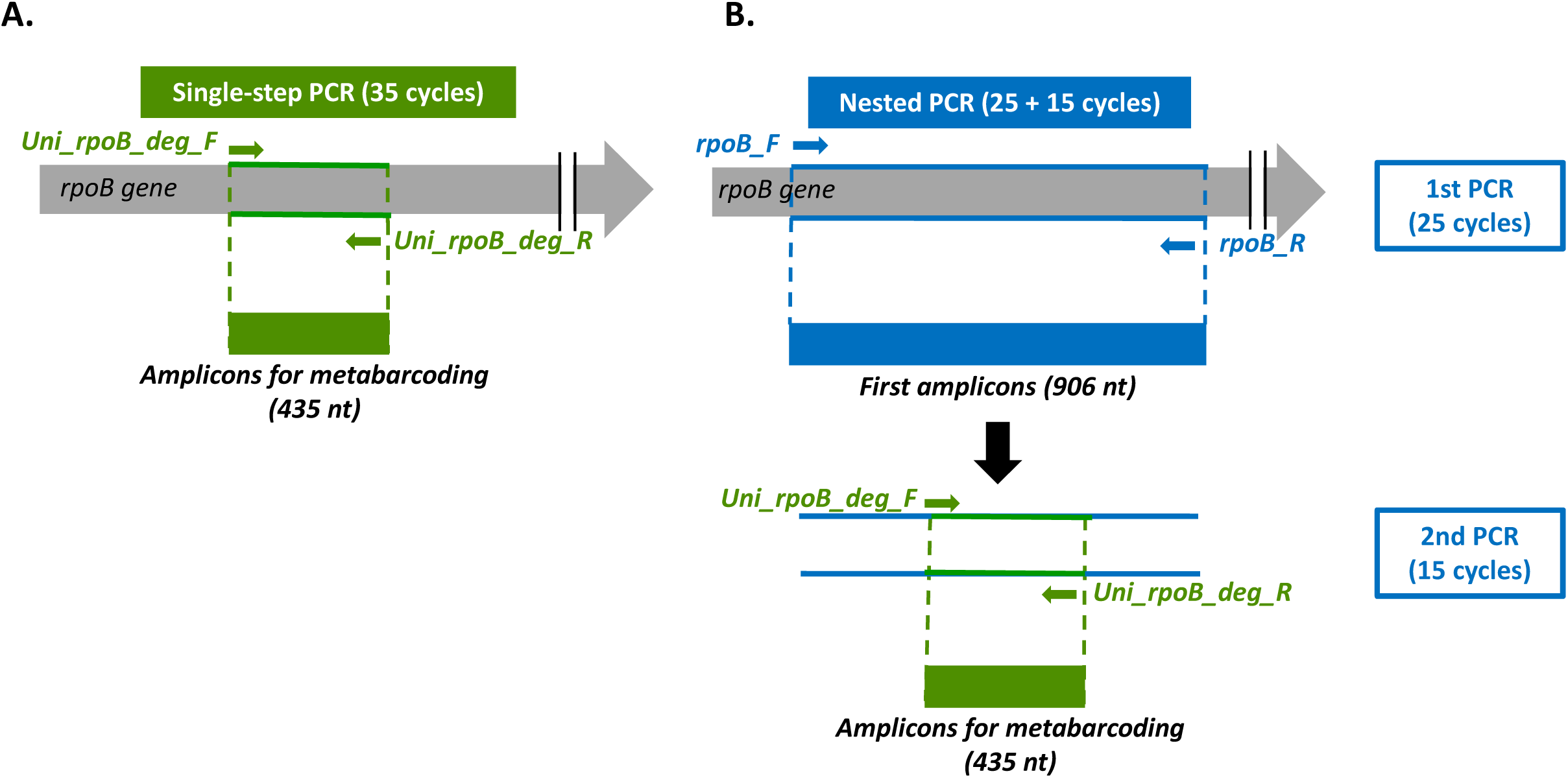
Protocol for the amplification of a region of the *rpoB* gene by **(A)** single-step PCR and **(B)** nested PCR strategies. The positions of the primers are indicated by green arrows (Uni_*rpoB*_deg_F/R primers) and blue arrows (*rpoB*_F/R primers). *nt*, nucleotides.

For the single-step *rpoB* PCR, we used the existing Univ_*rpoB*_deg_F (5’ - *TCG TCG GCA GCG TCA GAT GTG TAT AAG AGA CAG N* **GGYTWYGAAGTNCGHGACGTDCA** - 3’) and Univ_*rpoB*_deg_R (*5’ - GTC TCG TGG GCT CGG AGA TGT GTA TAA GAG ACA GN* **T GACGYTGCATGTTBGMRCCCATMA** - 3’) primers, which generate a 435 bp *rpoB* amplicon. These primers include a 5’ nucleotide overhang with adapters for Illumina library indexing, indicated by the bases in italics in the primer sequences above. The conditions for the single-step PCR *rpoB* were as previously described [5]. Briefly, 35 cycles of amplification were performed in a Bio-Rad thermocycler with 1 to 50 ng genomic DNA, the high-fidelity iProof™ DNA Polymerase (Bio-Rad), and an annealing temperature of 57⍰°C.

For the nested *rpoB* PCR, we designed degenerate consensus outer *rpoB* primers — *rpoB*_F (forward primer 5’ - **CAGYTDTCNCARTTYATGGAYCA**-3’) and *rpoB*_R (reverse primer 5’ - **AGTTRTARCCDTYCCANGKCAT** - 3’) — based on ClustalW alignments (http://multalin.toulouse.inra.fr/multalin/) of *rpoB* gene sequences from bacterial reference genomes representing a broad taxonomic diversity of eubacteria. The binding sites of the selected primers correspond to *Escherichia coli* K12 nucleotide positions 1528 to 2434, resulting in the amplification of 906-nucleotide portion of the *rpoB* gene. The specificity of the *rpoB* primers was checked *in silico* as previously described [5]. The first round of amplification in the nested PCR was performed with the *rpoB_F/R* primers and the GoTaq DNA Polymerase (Promega) in a Bio-Rad thermocycler, with 1 to 50 ng genomic DNA, in accordance with the enzyme manufacturer’s instructions. Twenty-five amplification cycles were performed (annealing temperature of 50 °C and annealing time of 45 s), generating an amplicon of 906 bp. The second round of amplification in the nested PCR was performed with 1 µL of the product of the first round of PCR, with the Univ_*rpoB*_deg_F/R primers incorporating Illumina adapters (see above) and the high-fidelity iProof™ DNA Polymerase (Bio-Rad). The same protocol was followed as for the single-step PCR (annealing temperature of 57 °C and annealing time of 45 s), except that the number of amplification cycles was reduced to 15.

The presence of contaminating DNA was assessed in each PCR run by including negative controls with sterile ultrapure water as the template. The amounts of amplicon DNA and the size of the amplicons were systematically analyzed by agarose gel electrophoresis.

All Illumina-indexed amplicons of the appropriate size on agarose gels were purified, multiplexed, and sequenced at the Genseq platform (University of Montpellier, France). The amplicons were purified with magnetic beads (Clean PCR, Proteigene, France) and a new round of PCR was performed in a total volume of 18 µL (5 µL of product from the first round of PCR, 9 µL Phusion® High-Fidelity PCR Master Mix, NEB, France, 2 µL index adapter I5, 2 µL index adapter I7). The cycling conditions were as follows: 95°C for 3 minutes, then 10 cycles of 95°C for 30 seconds, 55°C for 30 seconds, 72°C for 30 seconds, and a final elongation phase for 5 minutes at 72°C. We used a set of 384 index pairs based on the IDT’s unique dual index (UD) set to mark all samples and multiplex them in a single MiSeq run. The final PCR products were purified with magnetic beads, multiplexed and sequenced in pairs on an Illumina MiSeq sequencer with the MiSeq v3 reagent kit (600 cycles; Illumina) according to the manufacturer’s instructions. FastQ files were generated with bcltofastq at the end of the run. The quality of the raw data was assessed with a module developed by the MBB platform (Montpellier Bioinformatics Biodiversity platform supported by the LabEx CeMEB an ANR “Investissements d’avenir” program (ANR-10-LABX-04-01)) running the multiqc program [13].

The raw sequence data can be downloaded from http://www.ebi.ac.uk/ena/ (accession numbers: PRJEB81812)

### Bioinformatic processing of sequence data

The sequenced reads were analyzed with FROGS v4.1.0 software [14]. We successively used the pre-processing, clustering_swarm, remove_chimera, cluster_filter, taxonomic_affiliation tools with the parameters shown in Table S1.

### Update of the *rpoB* database

We created an updated *rpoB* sequence database by downloading the genomes with the genome_updater tool (https://github.com/pirovc/genome_updater). With this tool, we were able to create a RefSeq database [15] containing only complete genomes, complete chromosomes and protein sequences. We obtained protein sequences from 46,711 bacteria. We then used FetchMGS (https://github.com/motu-tool/fetchMGs.pl) [16] to extract DNA and protein sequence files for *rpoB*. We obtained 47,069 *rpoB* nucleic acid sequences from 46,298 bacteria. The *rpoB* database is available from the FROGS website: https://genoweb.toulouse.inrae.fr/frogs_databanks/assignation/rpoB/.

### *In silico* PCR analysis for evaluating the universality of *rpoB* primers

We used OBITools suite v1.2.11 [17] to evaluate the universality of the primers. The results were first formatted manually to make the identifiers unique and compatible with EcoPCR v1.0.1 [18], the *in silico* PCR tool. We then used *obiconvert* to construct a database that could be used by EcoPCR. We obtained three different types of amplicons targeting *rpoB*, by performing different runs of EcoPCR using: (i) the Univ_*rpoB*_deg_F and Univ_*rpoB*_deg_R primers (see above for the primer sequences) to obtain typical metabarcoding amplicons [5], (ii) the nested PCR primers to perform two successive PCR, one with *rpoB*_F and *rpoB*_R (see above) and the other based on previous output sequences with the Univ_*rpoB*_deg_F and Univ_*rpoB*_deg_R primers. For comparisons of taxonomic specificity, we compared only species to which these markers (amp_direct_*rpoB*, amp_nested_primary_*rpoB*, amp_nested_*rpoB*) were common. In total, 47,069 genomes were sufficiently complete to contain the *rpoB* gene.

The bacterial diversity of the amplicons obtained is displayed with the krona tool [19]. All these steps and scripts are detailed on github: https://github.com/geraldinepascal/RPOB_paper/tree/main/pcr_in_silico_script.

### Bacterial community and statistical analyses

ASV diversity and statistical analyses were performed with the R packages Phyloseq [20], Vegan [21], Microbiota Process [22] and Ampvis2 [23], as previously described [9]. Two custom-written scripts were used to generate plots to display diversity by taxonomy rank: add_multiaffi_to_abd_table.py (https://github.com/geraldinepascal/RPOB_paper/blob/main/FROGS_analysis_results/scripts/add_multiaffi_to_abd_table.py) and plot_taxo_ranks.py (https://github.com/geraldinepascal/RPOB_paper/blob/main/FROGS_analysis_results/scripts/plot_taxo_ranks.py).

## Results

### Design of single-step and nested ***rpoB*** PCR strategies for metabarcoding

The single-step and nested PCR strategies for the amplification of the *rpoB* gene are illustrated in Fig. 1. For the single-step PCR strategy (Fig. 1A), the region used for metabarcoding is directly amplified (35 cycles) with the Uni_*rpoB*_deg_F/R inner primers containing Illumina adapters [5] to yield a 435 bp amplicon. For the nested PCR strategy (Fig. 1B), we first designed the “*rpoB*_F/R” outer primers to bind to conserved regions of the *rpoB* gene encompassing the metabarcoding target region and generating a 906 bp amplicon. This amplicon was then subjected to a second amplification reaction with the inner primers incorporating Illumina adapters (Uni_*rpoB*_deg_F/R primers), generating a 435 bp amplicon. The first PCR should enrich the substrate for the second reaction in the *rpoB* target, thereby increasing the efficiency of the second round of PCR. We optimized the number of amplification cycles, using genomic DNA from biological samples available in the laboratory: 25 cycles for the first PCR and 15 for the second (data not shown). This optimization minimized the total number of cycles while ensuring a robust signal for Illumina sequencing and no detectable signal in the PCR negative control.

### *In silico* evaluation of the taxonomic coverage of *rpoB* primers

We checked the universality of the outer and inner *rpoB* primers *in silico* against an *rpoB* database with 47,069 entries, this database having been updated in 2024 (https://web-genobioinfo.toulouse.inrae.fr/frogs_databanks/assignation/rpoB/). The single-step and nested strategies had the potential to amplify 72% and 68%, respectively, of the 47,069 sequences stored in the *rpoB* database (Fig. 2). A Krona representation identified the bacterial taxa potentially amplified by the *rpoB* primers in the single-step PCR strategy (Fig. 2A) and after the first (Fig. S1) and second rounds (Fig. 2B) of the nested PCR strategy. The outer primers used in the first PCR of the nested approach had a high degree of taxonomic coverage, encompassing over 88% of all *rpoB* sequences, accounting for the similar levels of taxonomic coverage achieved with the single-step and nested strategies. The Pseudomonadota and most of the Bacillota, excluding the *Staphylococcaceae*, were well covered by the successive use of outer *rpoB* primers in the first PCR and inner *rpoB* primers in the second PCR. By contrast, the Actinomycetota and Bacteroidota were poorly covered by both the single-step and nested PCR strategies. For more details on taxonomic coverage, an interactive link is available from https://github.com/geraldinepascal/RPOB_paper/tree/main/krona_figure.

**Figure 2.**
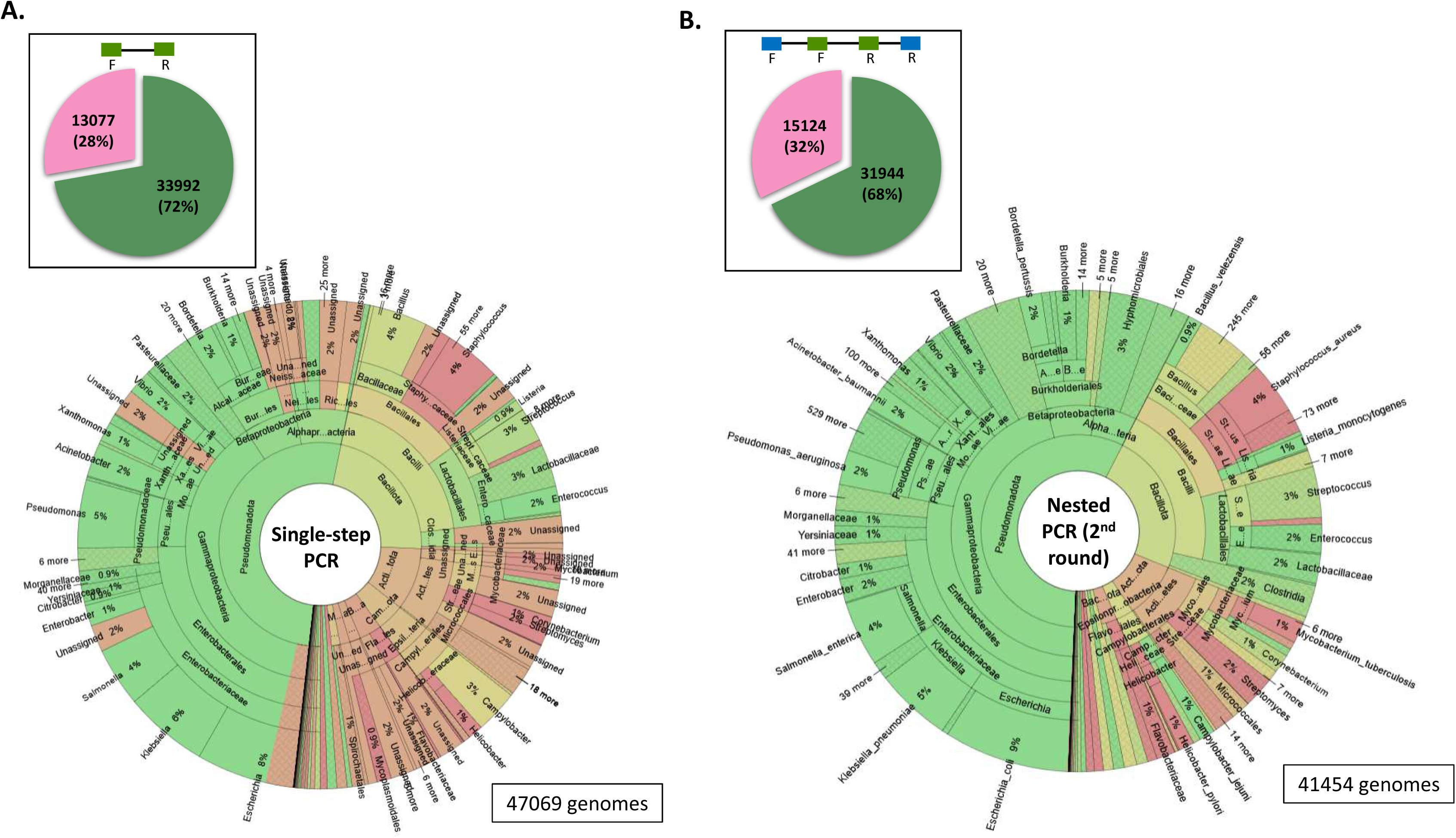
Coverage of bacterial taxonomic diversity, evaluated with the EcoPCR v1.0.1 *in silico* tool for **(A)** the single-step *rpoB* PCR strategy and **(B)** the nested *rpoB* PCR strategy. The *in silico* PCR was performed on the 47,069 *rpoB* sequences stored in the *rpoB* database. The pie charts illustrate the number of sequences amplified with two mismatches allowed (green) and the number of unretrieved sequences (pink) from the 47,069 sequences. Below, the Krona representations display the major taxa potentially amplified by the *rpoB* marker (green) in the single-step and nested PCR strategies. Taxa that are not amplified are shown in pink. For more details on the taxa covered by the two PCR strategies, all Krona visualizations are accessible via the interactive link: https://github.com/geraldinepascal/RPOB_paper/tree/main/krona_figure.

### Experimental comparison of the single-step and nested *rpoB* PCR strategies with different mock samples

We compared the sensitivity and reliability of nested and single-step PCR strategies on two commercial mock samples — Mock_8sp and Mock_8sp_log (see Table 1 for composition) — containing DNA from eight bacterial species, non-diluted (ND) or diluted 1/10 or 1/100. After amplification by PCR, gel electrophoresis of the *rpoB* amplicons showed that the nested PCR strategy resulted in successful amplification, even at dilutions as high as 1:100, for both mock samples (Fig. 3A and 3B). By contrast, with the single-step PCR strategy, amplification was possible only up to a dilution of 1/10 dilution for Mock_8sp (Fig. 3A) and only with the undiluted sample for the Mock_8sp_log (Fig. 3B). The nested PCR strategy was, therefore, more sensitive than the direct amplification approach. No amplification was observed from the negative control, regardless of the strategy used.

**Figure 3.**
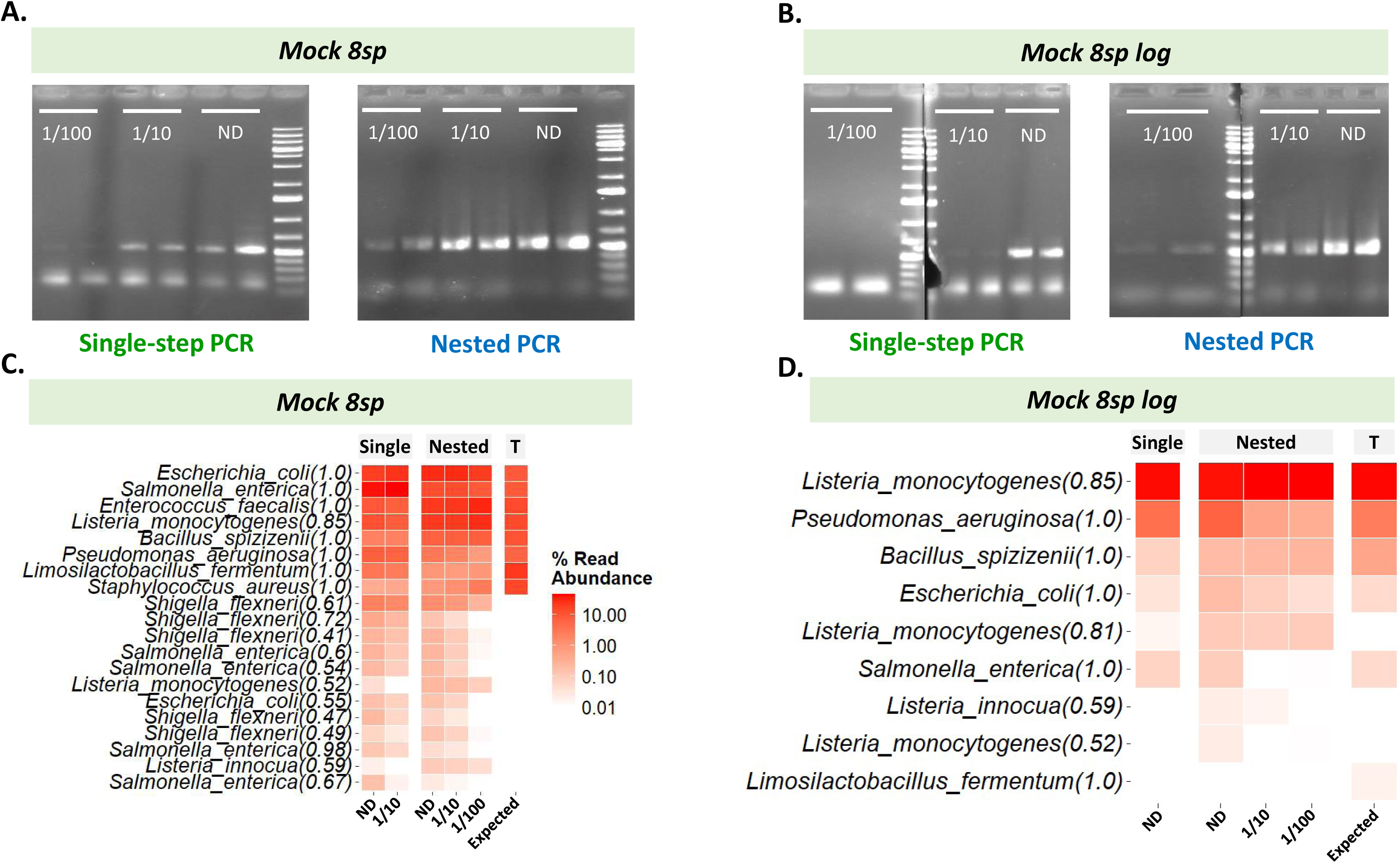
Comparison of single-step and nested *rpoB* PCR strategies with commercial mock communities used as DNA matrices to assess amplification sensitivity (**A** and **B**) and the ability to determine taxonomic composition by metabarcoding (**B** and **C**). Panels **A** and **B** show agarose gel electrophoresis of *rpoB* amplicons from mock_8sp and mock_8sp_log samples, respectively, amplified by single-step or nested PCR strategies. The dilutions of the mock samples are indicated on the agarose gel, below the wells. For each dilution, two technical replicates are shown. ND: not diluted. Panels **C** and **D** show heatmaps for the bacterial taxonomic composition obtained by *rpoB* metabarcoding of the mock_8sp and mock_8sp_log samples, respectively (only samples with successful amplification were sequenced). The columns ‘Single’ and ‘Nested’ correspond to the single-step and nested *rpoB* PCR strategies, and the column ‘T’ indicates the expected composition of the mocks. The dilutions of the mock samples are indicated at the bottom of the heatmap. The 20 most abundant ASVs across samples, assigned at species level, are listed on the left. Each species name is followed by an RDP bootstrap confidence score, indicating the confidence level of the taxonomic assignment. The relative read abundance is represented by a gradient of red hues.

We then sequenced the amplicons visible on agarose gels with Illumina MiSeq technology. The number of reads per sample and the ASV composition are detailed in Table S2; rarefaction curves are shown in Fig. S2. The relative abundance of ASVs by taxonomic rank clearly confirmed that the *rpoB* primers provided a high degree of taxonomic resolution, with 100% of taxa assigned to species level in the mock samples (Fig. S3). The bacterial compositions obtained for Mock_8sp after the single-step and nested amplification strategies closely resembled the theoretical composition of the mock sample (Fig. 3C and 3D). However, several additional sequences were assigned to the species *Shigella flexneri* in Mock_8sp and *Listeria innocua* in both mock samples, regardless of the strategy used. These additional assignments probably correspond to artifactual sequence variants, attributable to the close phylogenetic relationship between the genera *Shigella* and *Escherichia*, or between *L. innocua* and *L. monocytogenes*. However, neither *E. faecalis* nor *S. aureus* was detected in Mock 8sp_log by either single-step or nested PCR, consistent with the low proportion of genomes attributable to these species in Mock 8sp_log, below 0.1%, the threshold generally required for signal detection by an amplicon metabarcoding approach. We then compared the relative abundance of taxa at species level obtained for the Mock_8sp samples with the single-step and nested *rpoB* PCR strategies (Fig. S4). Both PCR strategies overestimated the abundances of *E. coli, Enterococcus faecalis*, and *Listeria monocytogenes*, while significantly underestimating the abundances of *Limosilactobacillus fermentum* and *Staphylococcus aureus*. This result is consistent with the *in silico* analysis of the taxonomic diversity targeted by the *rpoB* primers, which revealed limited coverage for certain bacterial groups within the Bacillota (see above). The single-step PCR strategy significantly overestimated the abundance of *Salmonella enterica,* whereas the nested PCR strategy did not (Fig. S4).

Overall, our results demonstrate that the nested PCR strategy provides taxonomic assignment profiles similar to those generated by the single-step method, but with a greater sensitivity and more efficient amplification for samples containing a low concentration of the target DNA.

### Application of the nested *rpoB* strategy to characterization of the microbiota of insect samples

We experimentally validated the nested PCR strategy on an insect, *Spodoptera frugiperda*, sampled in Senegal (near Velingara). We tested our methodology on two types of biological matrix: (i) DNA extracted from the oral secretion (OS) of larvae, a biological fluid composed of saliva, the gut microbiota, and a food bolus [24], which contains only very low concentrations (< 4 µg/mL) of bacterial DNA, and (ii) DNA extracted from demi-larvae (caterpillars), in which the bacterial DNA is embedded in a eukaryotic DNA matrix. The *rpoB* region was targeted by both the single-step and nested PCR strategies. PCR amplifications were performed on the six replicate samples of larval DNA (three stage L3, three stage L5), on DNA from the pooled OS sample, for which three PCR replicates were processed, and on the extraction control samples. The single-step *rpoB* PCR strategy generated no visible amplicon for either the OS or larval samples, which it was therefore impossible to sequence for metabarcoding. By contrast, the nested *rpoB* strategy successfully amplified all OS and larval samples, yielding sufficient DNA for metabarcoding. Importantly, no amplification was observed for the extraction or PCR negative controls following the two successive PCR rounds of the nested *rpoB* strategy.

All amplicons generated by the *rpoB* nested strategy were further subjected to Illumina MiSeq sequencing and statistical analysis. Details of the read count per sample and ASV composition are provided in Table S2, and rarefaction curves are provided in Fig. S5. The Pseudomonadota largely dominated in the pooled OS sample, accounting for three quarters of the observed diversity, with Bacillota accounting for the remaining quarter (Fig. 4A). Two species predominated (Fig. 4B): a species belonging to the genus *Ochrobactrum*, but with an insufficient RDP identification confidence threshold for reliable assignment to a species, and a species confidently identified (RDP bootstrap confidence of 1) as *E. casseliflavus*. There were slight variations between technical PCR replicates, highlighting the importance of such replicates for ensuring reliable and reproducible results for the metabarcoding of low-concentration DNA samples.

**Figure 4.**
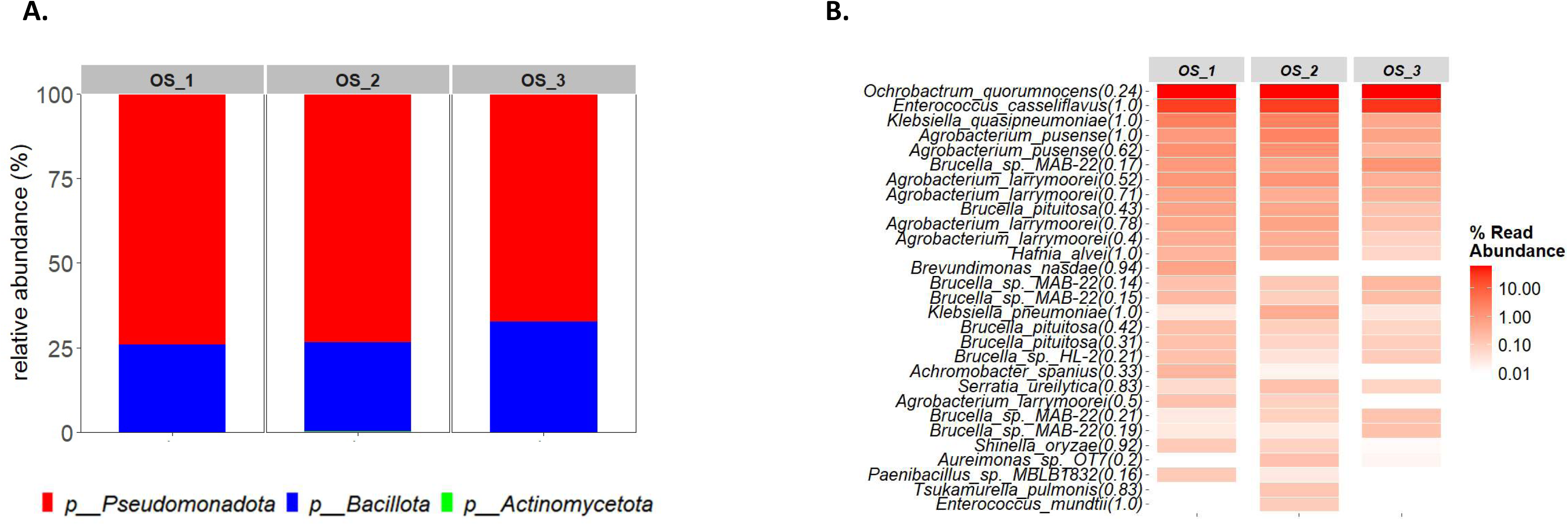
Composition of the bacterial community according to metabarcoding of oral secretion (OS) samples from *S. frugiperda* amplified by the nested *rpoB* PCR strategy. (***A***) At phylum level, the relative abundances are represented on bar plots, with sample names indicated at the top of the plots. (***B***) At species level, the 30 most abundant ASVs across samples are listed on the left of the heatmap. Each species name is followed by an RDP bootstrap confidence score, indicating the confidence level of the taxonomic assignment. The relative abundance (percentage) is represented by a gradient of red hues. Sample names, corresponding to the three PCR replicates, are indicated at the top of the heatmap columns.

For larval samples, the nested *rpoB* strategy revealed a predominance of Pseudomonadota in five of the six biological replicates, with only one L5-stage sample containing a majority of Bacillota (Fig. 5A). The nested *rpoB* strategy performed well for identification to species level (i.e., RDP assignment threshold generally > 0.8), making it possible to identify the species associated with *S. frugiperda* larvae with confidence (Fig. 5B). In particular, the metabarcoding analysis revealed the prevalence of *Klebsiella quasipneumoniae, Acinetobacter soli, E. casseliflavus, K. pneumoniae,* and *A. baumannii* in both L3-stage and L5-stage caterpillars. Interestingly, a few taxa, such as *E. casseliflavus and K. quasipneumoniae,* were common to both the larval and OS samples.

**Figure 5.**
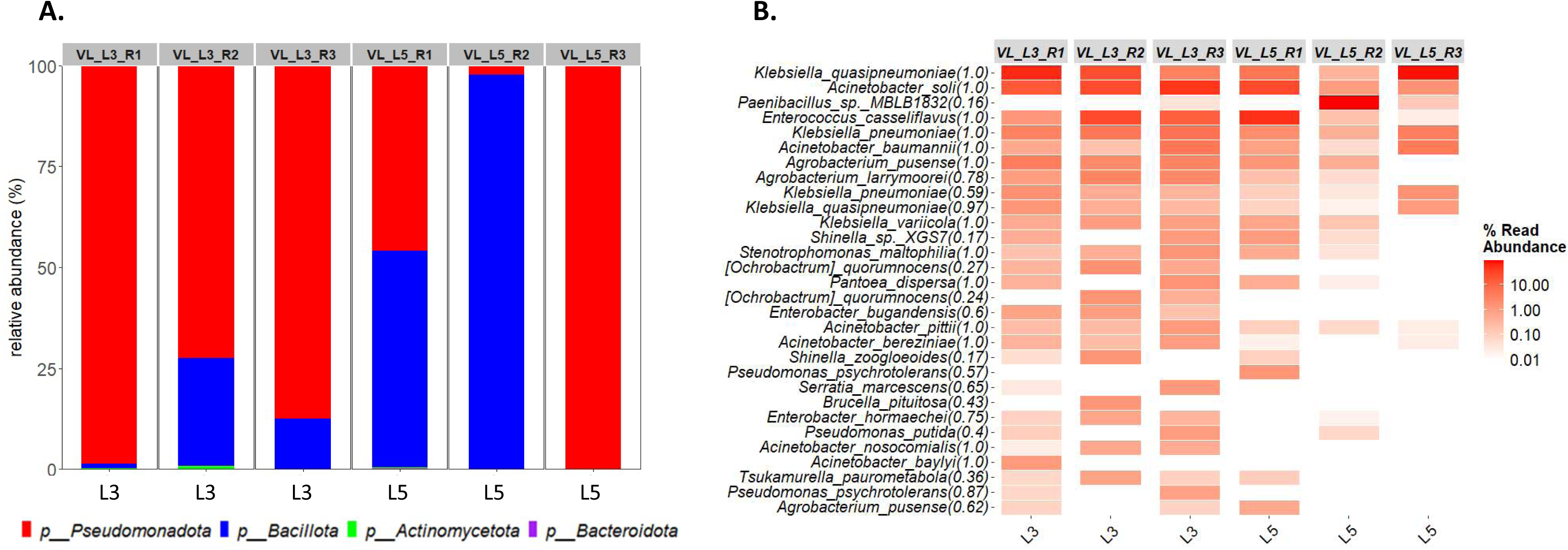
Composition of the bacterial community revealed by metabarcoding of *S. frugiperda* demi-larva samples subjected to amplification with the nested *rpoB* PCR strategy. (***A***) At phylum level, relative abundances are represented on bar plots, with sample names indicated at the top of the plots. At the bottom of the graph, L3 corresponds to the third developmental stage, and L5 corresponds to the fifth developmental stage of the caterpillar (***B***) At species level, the 30 most abundant ASVs across samples are listed on the left of the heatmap. Each species name is followed by an RDP bootstrap confidence score, indicating the confidence level of the taxonomic assignment. The relative abundance (percentage) is represented by a gradient of red hues. Sample names are indicated at the top of the heatmap columns.

## Discussion

Metabarcoding is a powerful tool for simultaneously identifying multiple taxa within a DNA sample. However, short-read metabarcoding studies, which typically rely on amplicons of variable regions of the 16S rRNA gene to characterize bacterial microbiota [25–27], lack the precision required for fine-scale taxonomic resolution [28, 29].

In this study, we focused on the *rpoB* marker due to its ability to provide species-level taxonomic assignments [5], providing a promising alternative to 16S rRNA-based approaches. However, the primers for *rpoB* have several weaknesses. Due to the lack of highly conserved regions in the *rpoB* gene, the primers contain multiple degenerate bases, which can make specific amplification difficult, particularly in matrices with low concentrations of bacterial gDNA and in host-associated samples, which generally contain much less prokaryotic than eukaryotic DNA. For example, even with 35 amplification cycles, PCR with *rpoB* primers fails to amplify the target efficiently in some complex host-associated microbiota, such as insect samples. Bivand and coll. [30] tried to improve the specificity of the primers for target binding by designing *rpoB* primers according to the dual priming oligonucleotide (DPO) principle, and coupling the amplification with Sanger sequencing to detect pathogenic taxa in clinical samples dominated by human DNA. This approach was effective for targeted detection but is not suitable for NGS-based metabarcoding.

In this study, we validated a nested PCR strategy to improve the amplification efficiency and sensitivity of *rpoB* primers for metabarcoding analysis. Unlike the standard single-step *rpoB* PCR, this nested PCR involves a first PCR to enrich the sample in the target DNA and a second PCR to amplify the *rpoB* fragments for metabarcoding. Nested PCR has long been known to increase PCR sensitivity [31] but, to our knowledge, this is the first time that this method has been used in a metabarcoding approach.

We first demonstrate the reliability of nested PCR with the *rpoB* marker for the accurate identification of bacterial species in different mock samples of known composition and variable complexity, and its ability to increase amplification sensitivity relative to the single-step PCR strategy. Some studies have reported that nested PCR has the drawback of potentially increasing bias through preferential amplification in the two successive PCR steps [32]. By optimizing the number of cycles between the first and second PCRs (40 cycles in total), we minimized the biases associated with nested PCR, ensuring robust results comparable to those obtained by single-step PCR (35 cycles in total).

We then compared the performance of the single-step and nested *rpoB* PCR strategies for describing the host-associated bacterial microbiota in two types of lepidopteran insect samples: (i) *S. frugiperda* larval oral secretions (OS), corresponding to a matrix containing low concentrations of bacterial DNA and, (ii) *S. frugiperda* larvae, corresponding to a eukaryotic matrix with low proportions of bacterial DNA. The single-step *rpoB* PCR failed to amplify the target, whereas the *rpoB* nested PCR strategy successfully amplified the target from all insect samples. The *rpoB* nested PCR strategy is, therefore, considerably more sensitive than the single-step *rpoB* PCR, for samples with low concentrations of DNA and samples in which eukaryotic DNA predominates over bacterial DNA.

However, our findings also reveal several limitations of the *rpoB* primers, such as their limited coverage of certain bacterial phyla, contrasting with the universality of 16S primers. Indeed, the *rpoB* primers cover only about 70% of bacterial diversity (essentially the phyla Pseudomonodota and Bacillota). For example, *rpoB* lacks sensitivity for the detection of Bacteroidota, members of which may be present in host-associated microbiota samples, such as insect samples. As a means of addressing this bias, we propose the design of a multi-marker approach using phylum-specific *rpoB* primer pairs to achieve an optimal coverage of bacterial diversity.

## Conclusion

In metabarcoding studies, combining a nested PCR strategy with the use of the *rpoB* marker appears to be a promising approach for describing the bacterial complexity and richness of insect-associated microbiota directly at species level. Our study highlights the benefits of nested PCR for efficiently amplifying prokaryotic DNA from eukaryotic hosts. The nested PCR *rpoB* metabarcoding method, with its species-level resolution and enhanced sensitivity, offers new opportunities for investigating the ecological roles of specific bacterial taxa in insect physiology, health, and environmental interactions. Finally, it would be of interest to extend this strategy to other host-associated microbiota samples. This study provides valuable insights for the growing scientific community exploring bacterial communities through metabarcoding across diverse ecosystems.

## List of abbreviations

DNA: deoxyribonucleic acid
OS: oral secretion
ASV: amplicon sequence variants
PCR: polymerase chain reaction
RDP Classifier: Ribosomal Database Project classifier

## Declarations

### Ethics approval and consent to participate

Not applicable

### Consent for publication

Not applicable

### Availability of data and material

The sequence data supporting the findings of this study have been deposited in the European Nucleotide Archive under primary accession code PRJEB81812.

### Authors’ contributions

L.L., S.G., J.C.O. conceived and designed the experiments. L.L, J.C.O. conducted the experiments. L.L., S.G., J.C.O. analyzed the data. J.C.O., G.A., J.M. and G.P. conducted the bioinformatics analysis and analyzed the data. T.B. organized and led the Senegal mission. LL and SY collected FAW larvae in Senegal. LL collected the OS in the field. A.C., N.N, S.G and J.C.O. supervised the research and contributed to the discussion of the results. L.L., S.G., J.C.O. wrote the manuscript. L.L., J.C.O., G.A., G.P. prepared the figures and supplementary files. All authors reviewed the manuscript.

## Acknowledgments

We would like to thank the Insect Quarantine Platform (PIQ), a member of the Vectopole Sud network, for providing the infrastructure required for experiments on insect pests. We would also like to thank Gaëtan Clabots, Raphaël Bousquet and Dylan Valenza for maintaining the FAW colonies at the DGIMI laboratory in Montpellier.

The metabarcoding data were produced with the GenSeq technical facilities at the “Institut des Sciences de l’Evolution” in Montpellier, with support from LabEx CeMEB, an ANR “Investissements d’avenir” program (ANR-10-LABX-04-01).

We would like to thank the Genotoul bioinformatics platform Toulouse Occitanie (Bioinfo Genotoul, https://doi.org/10.15454/1.5572369328961167E12) for providing assistance, computing and storage resources.

We would like to thank Maïmouna Cissoko and Nils Poulicard for their help during field sampling in Senegal and Victoria Mariotti for her help with bioinformatics analyses.

## Supplemental Materials

**Table S1.** FROGS parameters

**Table S2.** Detailed amplicon sequence variants (ASV) table for the samples analyzed

**Fig. S1.** Krona representations of the bacterial taxa potentially amplified by the outer primers used in the first round of PCR in the nested *rpoB* PCR strategy. The *in silico* EcoPCR v1.0.1 tool was used on the 47,069 *rpoB* sequences stored in the *rpoB* database, with two mismatches allowed. The Krona graph displays the major taxa potentially amplified by the outer *rpoB* primers (in green). Taxa that are not amplified are shown in pink. For more details on the taxa covered by the two PCR strategies, all Krona visualizations are accessible via the interactive link: https://github.com/geraldinepascal/RPOB_paper/tree/main/krona_figure.

**Fig. S2.** Rarefaction curves for the metabarcoding sequences of the different commercial mock samples amplified by the single-step or nested *rpoB* PCR strategy. For each sample, species richness is shown on the *y*-axis and the number of sequences is shown on the *x*-axis. The ggrare function was used to generate the rarefaction curves (https://rdrr.io/github/gauravsk/ranacapa/man/ggrare.html).

**Fig. S3.** Plot showing the relative abundance of ASVs at different taxonomic ranks for the various commercial mock samples following amplification by the single-step or nested *rpoB* PCR strategy. Two custom scripts were used to generate the plot: add_multiaffi_to_abd_table.py (https://github.com/geraldinepascal/RPOB_paper/blob/main/FROGS_analysis_results/scripts/add_multiaffi_to_abd_table.py) and plot_taxo_ranks.py (https://github.com/geraldinepascal/RPOB_paper/blob/main/FROGS_analysis_results/scripts/plot_taxo_ranks.py).

**Fig. S4.** Histogram plot of bacterial composition showing the relative abundances of ASVs at species level in the non-diluted commercial mock_8sp sample, amplified by the single-step or nested *rpoB* PCR strategy. Two technical replicates (rep1 and rep2) were performed, with the last bar indicating the expected composition.

**Fig. S5.** Rarefaction curves for the metabarcoding sequences of *S. frugiperda* OS samples (***panel A***) and *S. frugiperda* demi-larva samples (***panel B***) subjected to amplification by the single-step or nested *rpoB* PCR strategy. The OS samples correspond to three independent PCR replicates. The larval samples correspond to several biological replicates: L3 is the third developmental stage, and L5 is the fifth developmental stage of the caterpillar. For each sample, species richness is displayed on the *y*-axis and the number of sequences on the *x*-axis. The ggrare function was used to generate the rarefaction curves (https://rdrr.io/github/gauravsk/ranacapa/man/ggrare.html).

