## Supplementary material for "Nested PCR to optimize *rpoB* metabarcoding for low-concentration and host-associated bacterial DNA": Fig. S1

### Slide 1
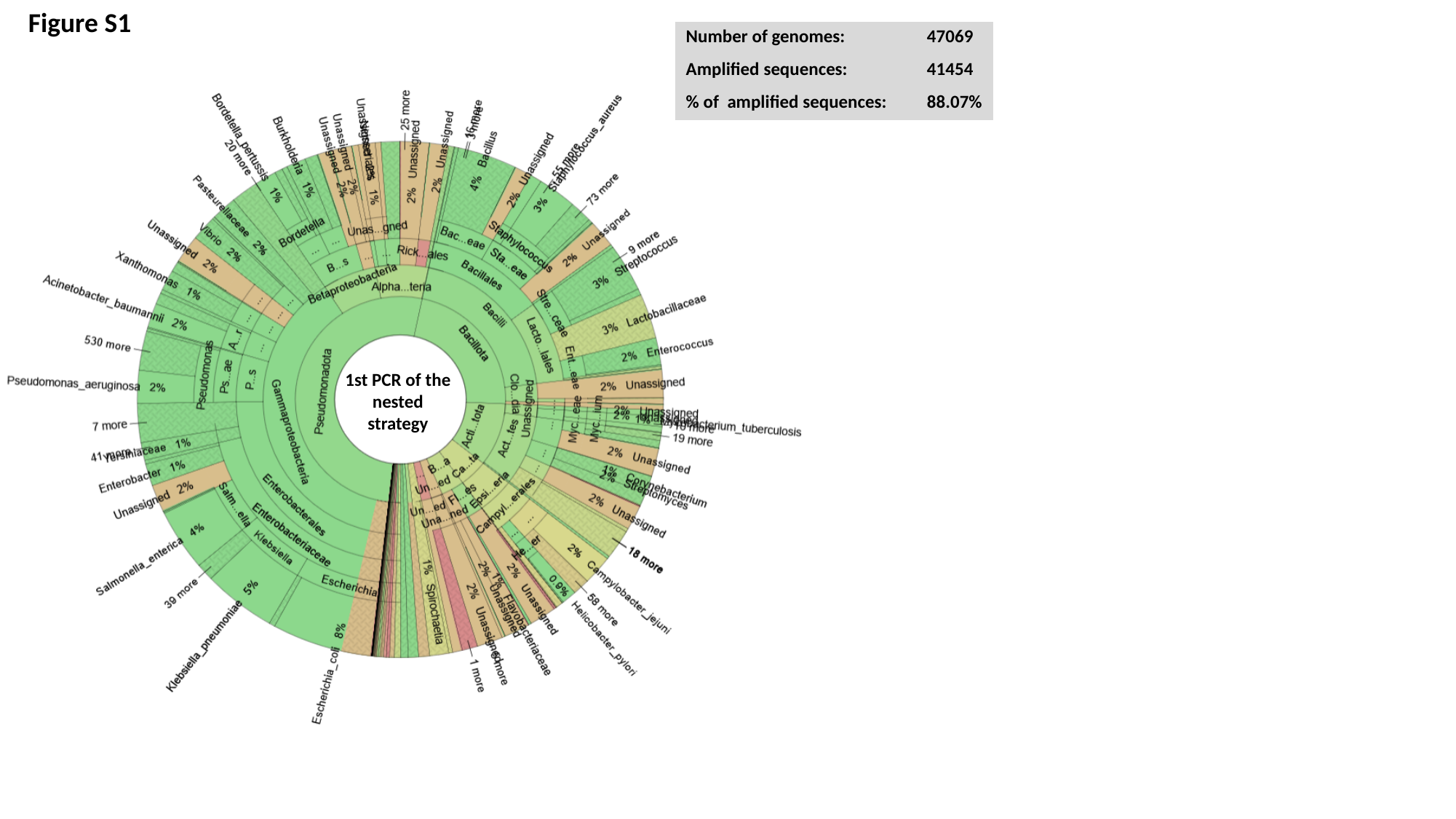

Figure S1
| Number of genomes: | 47069 |
| --- | --- |
| Amplified sequences: | 41454 |
| % of amplified sequences: | 88.07% |
1st PCR of the nested strategy
