## Supplementary material for "Nested PCR to optimize *rpoB* metabarcoding for low-concentration and host-associated bacterial DNA": Fig. S2

### Slide 1
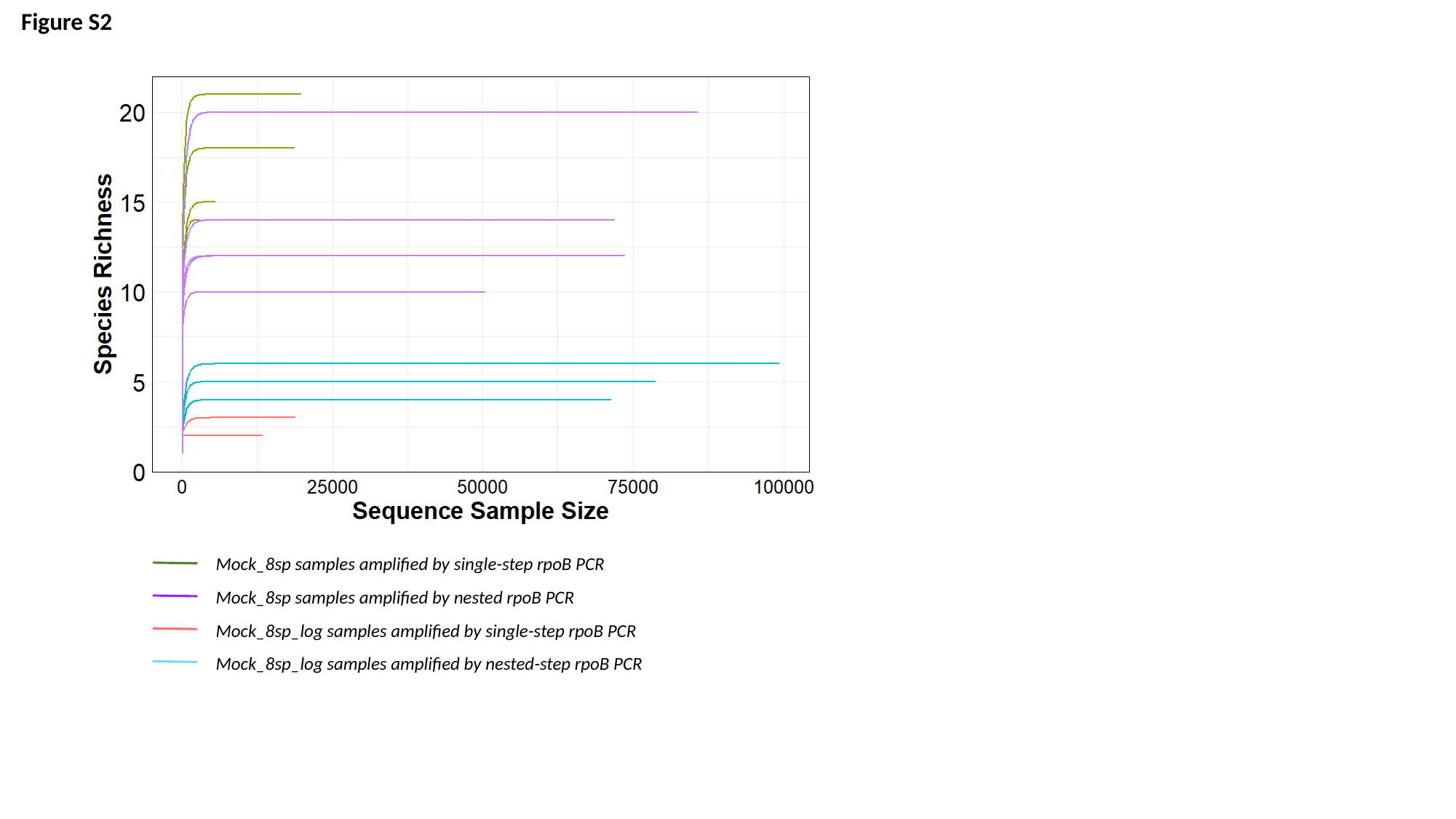

Figure S2
Mock_8sp samples amplified by single-step rpoB PCR
Mock_8sp samples amplified by nested rpoB PCR
Mock_8sp_log samples amplified by single-step rpoB PCR
Mock_8sp_log samples amplified by nested-step rpoB PCR
