## Supplementary figures and images for "Nested PCR to optimize *rpoB* metabarcoding for low-concentration and host-associated bacterial DNA"

### Fig. S3

#### Slide 1
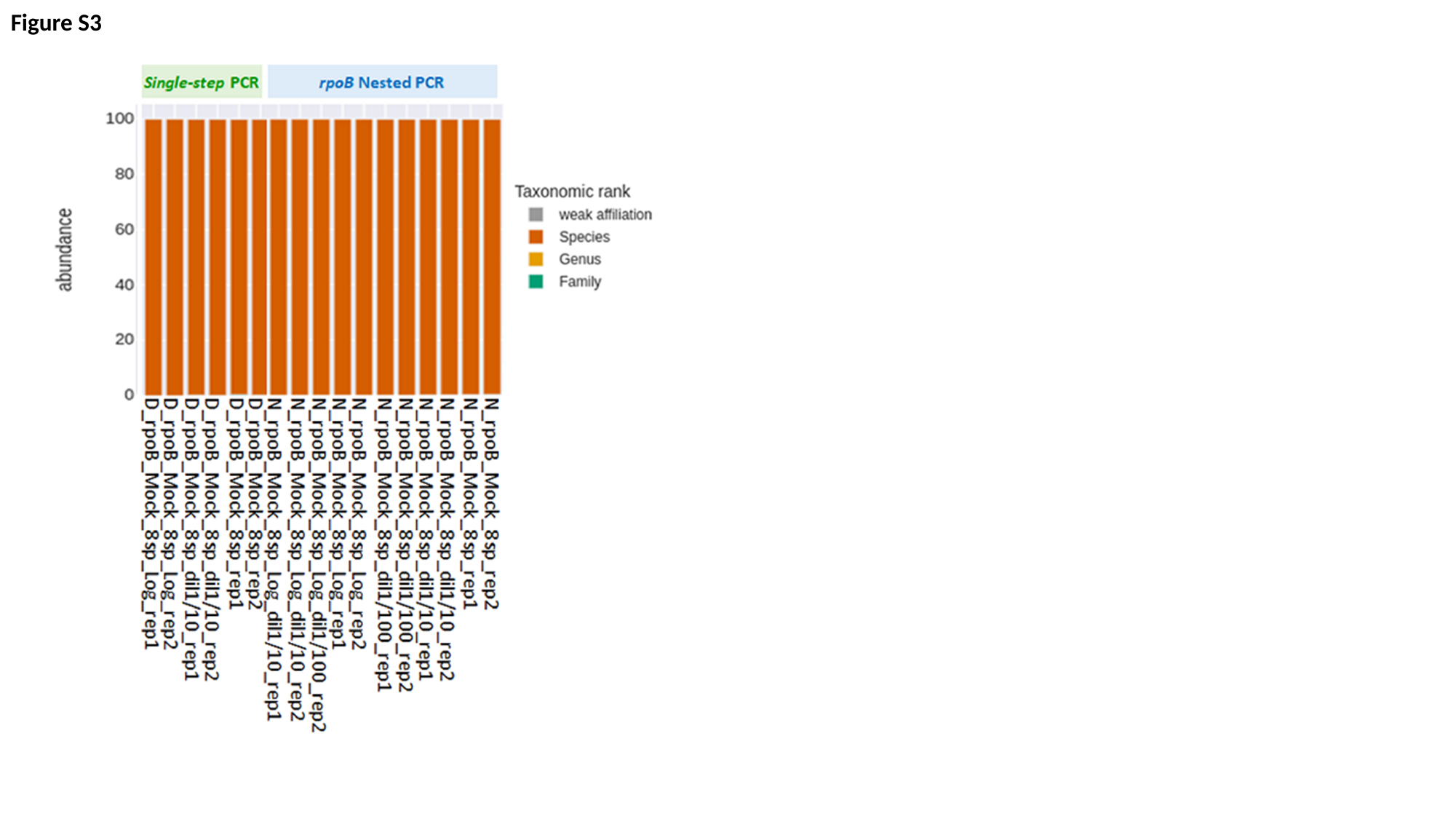

Figure S3

### Fig. S4

#### Slide 1
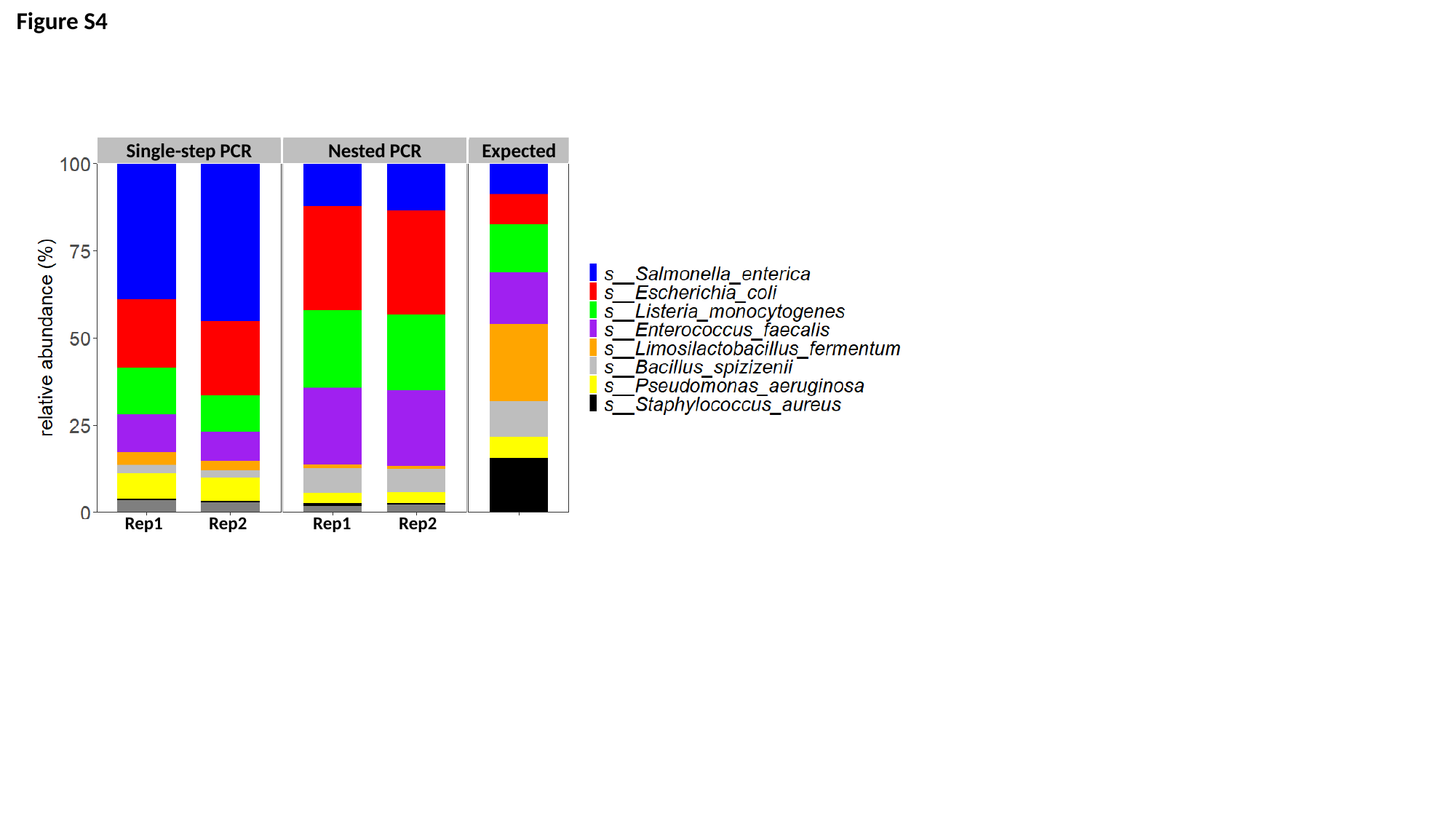

Figure S4
Single-step PCR
Nested PCR
Expected
Rep1
Rep2
Rep1
Rep2

### Fig. S5

#### Slide 1
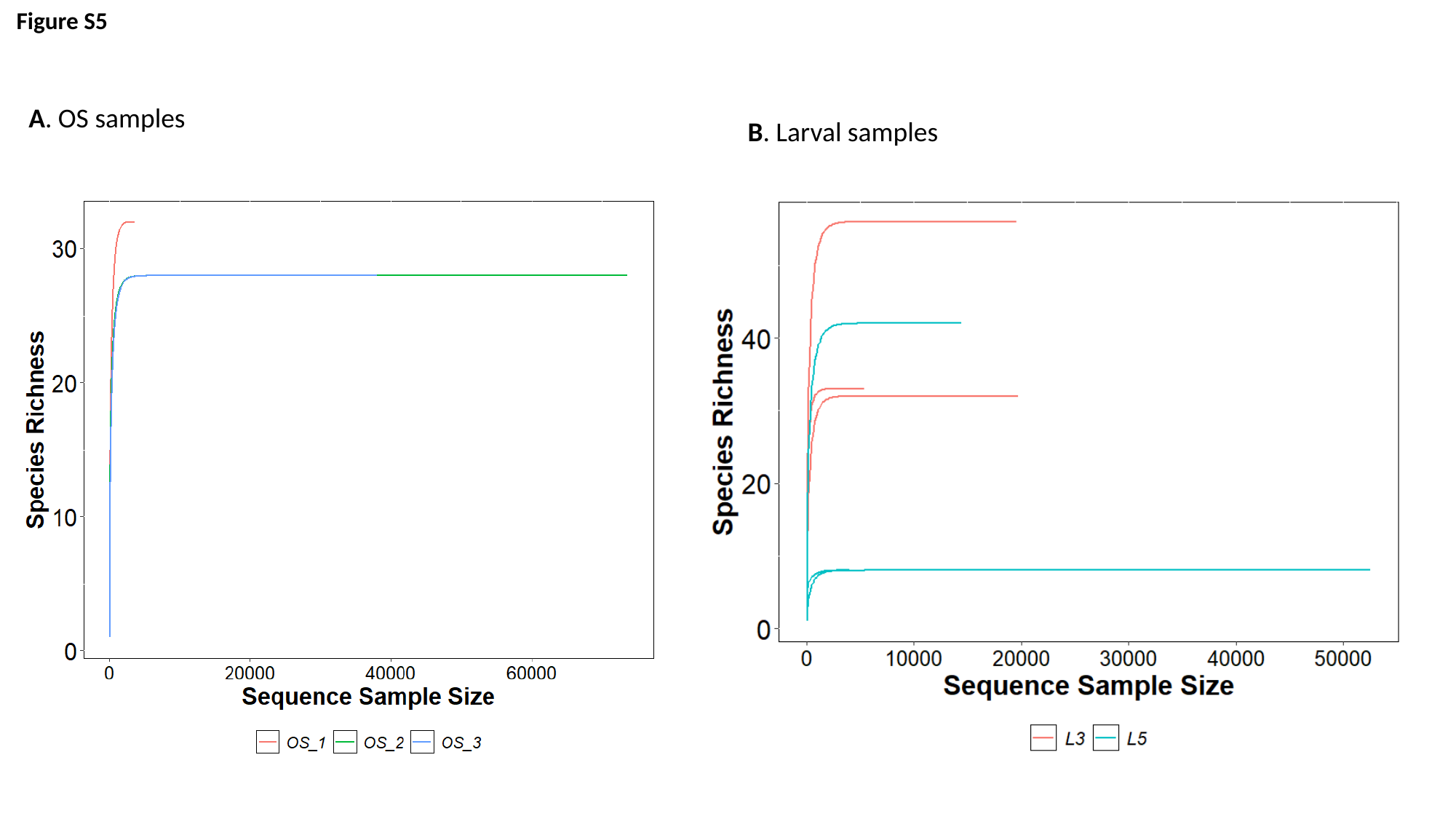

Figure S5
A. OS samples
B. Larval samples
