## Supplementary material for "Nested PCR to optimize *rpoB* metabarcoding for low-concentration and host-associated bacterial DNA": Table S1

**Table S1** : FROGS 4.1.0 parameters used for this study for *rpoB* amplicons from larvae, oral secretion and mock samples.

| **Tool** | ***rpoB*** |
| --- | --- |
| **preprocessing** | Illumina  --min-amplicon-size 300 --max-amplicon-size 590 –five-prim-primer GGYTWYGAAGTNCGHGACGTDCA --three-prim-primer TKATGGGYKCVAACATGCARCGTCA  --R1-size 300 --R2-size 300 |
| **clustering** | --fastidious --distance 1 |
| **remove_chimera** | default |
| **cluster_filters** | --min-abundance 0.00005 |
| **taxonomic_affiliation** | --rdp  --reference rpoB_bacteria_NCBI_refseq_genome_complete_and_chromosome_20240707.fasta |
